# The Moran coalescent in a discrete one-dimensional spatial model

**DOI:** 10.1101/184705

**Authors:** Max Shpak, Jie Lu, Jeffrey P. Townsend

**Author notes:** Corresponding author: Max Shpak, St. David’s Medical Center, 1015 E. 32nd St, Suite 414, Austin TX 78705.

## Abstract

Among many organisms, offspring are constrained to occur at sites adjacent to their parents. This applies to plants and animals with limited dispersal ability, to colonies of microbes in biofilms, and to other genetically heterogeneous aggregates of cells, such as cancerous tumors. The spatial structure of such populations leads to greater relatedness among proximate individuals while increasing the genetic divergence between distant individuals. In this study, we analyze a Moran coalescent in a one-dimensional spatial model where a randomly selected individual dies and is replaced by the progeny of an adjacent neighbor in every generation. We derive a recursive system of equations using the spatial distance among haplotypes as a state variable to compute coalescent probabilities and coalescent times. The coalescent probabilities near the branch termini are smaller than in the unstructured Moran model (except for *t* = 1, where they are equal), corresponding to longer branch lengths and greater expected pairwise coalescent times. The lower terminal coalescent probabilities result from a spatial separation of lineages, i.e. a coalescent event between a haplotype and its neighbor in one spatial direction at time *t* cannot co-occur with a coalescent event with a haplotype in the opposite direction at *t* + 1. The concomitant increased pairwise genetic distance among randomly sampled haplotypes in spatially constrained populations could lead to incorrect inferences of recent diversifying selection or of population bottlenecks when analyzed using an unconstrained coalescent model as a null hypothesis.

## 1 Introduction

Most of the statistical tests used to identify the population genetic signatures of natural selection or of population dynamics use neutral models of genetic variation as null hypotheses for the distribution of observed genetic variation. For example, under a neutral infinite sites model of evolution, the expected pairwise genetic distance between two randomly selected haplotypes is the product of the mutation rate and the expected coalescent time *T_ij_* among pairs *i, j*, i.e.

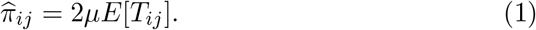

where the estimated mean pairwise genetic distance in a sample of size *n* is

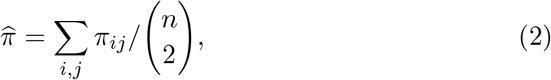

for Hamming distance *π_ij_* between haplotypes *i* and *j*. Following Tajima (1989), 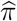 is an estimator for the diploid population mutation rate *θ* = 4*Nµ* (for a population of *N* haplotypes, *θ* = 2*Nµ*).

Estimators for *θ* are the basis for several statistical tests that distinguish selection from neutral evolution, or, alternatively, distinguish equilibrium population dynamics from recent changes in population size (Nielsen, 2001). Consequently, population characteristics that decrease or increase *E*[*T_ij_*] relative to the neutral model can lead to mistaken inferences of nonneutral evolution or non-equilibrium population dynamics. For example, in the Tajima’s D test, a distribution of coalescent times with deeper branches than those predicted under a neutral equilibrium model lead to inferences of either balancing selection or of recent population bottlenecks that disproportionately eliminate all but the most common alleles with the deepest branches on the coalescent tree.

The standard coalescent model (Kingman 1982, Hudson 1988, Hein et al 2005, Wakeley 2009) assumes a fully mixed, unstructured population undergoing Fisher-Wright genetic drift. In most natural populations, genetic variation is spatially partitioned due to the proximate co-occurrence of related individuals; this isolation by distance is especially true in populations of sessile or nearly sessile organisms with limited dispersal ability. Examples include vegetatively reproducing plants and animals, microbes growing on biofilms, cancer cells proliferating as tumors—indeed, any spatially constrained biological system where progeny remain proximate to their birthplace.

The effects of spatial structure on coalescent times have been previously investigated, especially the structured coalescent in island models (e.g. Slatkin 1987) that separate coalescent times within demes from the waiting time to coalesce among demes via migration. For sufficiently small migration rates, this scenario can be modeled via a separation of timescales (Moehl 1998, 2000, Wakeley and Aliacar 2001, Wilkins 2004, Shpak et al 2010). The coalescent for Fisher-Wright genetic drift in a continuous spatial model with a diffusion model of migration was investigated in Wilkins and Wakeley (2002), who showed that the separation of timescales also occurs in a continuous spatial model if the population size and physical distance is an order of magnitude greater than the per-generation migration distance (see also Kelleher et al 2014, Joseph et al 2016).

In this paper, we consider a discrete model of linear spatial structure in the liming case where all progeny are constrained to appear at a position one unit of distance away from their parents. Using a Moran model for birth and death, we will derive a recursive system of equations to numerically calculate the coalescent probabilities in this discrete linear spatial model. Our approach creates a foundation upon which more general models of coalescent processes in discrete spatially structured populations can be built, as well as providing heuristic insight into the effect of spatial structure on coalescent times and probabilities.

## 2 A discrete one-dimensional Moran model

Consider a population of *N* haploid individual organisms, located on a line, where death and reproduction follow a modified Moran coalescent process (Moran 1958, see also Nowak 2006 Ch. 6, Etheridge and Griffiths 2009, Etheridge et al 2010). In every generation, a randomly selected individual *a_i_* at position *i* = 1…*N* on the line is selected to die. This individual can only be replaced by the progeny of an immediate neighbor at positions *i* + 1 or *i* – 1; the specific parent is randomly selected between these two neighbors with equal probability. In backward time, this specification means that in the previous generation, an individual haplotype at position *i* has occupied the same position and persisted, or it has been replaced by a descendant of an adjacent neighbor.

We use *a_i_*(*t*) to denote the “ancestor” of individual *a_i_* at exactly *t* generations ago, whether *a_i_*(*t*) represents a different, ancestral haplotype or simply persistence of the same individual over these time intervals. Note that *a_i_*(*t*) is not necessarily or typically located in position *i* at generation *t* ≠ 0, the *i* index denotes only the position of the lineage when it was sampled at *t* = 0. By abuse of notation, we use *a_i_* in place of *a_i_* (0), and we will use the expression *P_t_*(*i,j*) to represent the probability that haplotypes *a_i_,a_j_* coalesce at time *t*.

Because each generation is characterized by a single death and a single replacement, there are *N* – 1 possible coalescent events for a population on a bounded line, because haplotypes that are located “internally” can coalesce with either a left or right neighbor, while those at endpoints can coalesce with only a single neighbor. If individuals are arranged on a circle, there are *N* possible coalescent events in every generation, because each haplotype can coalesce with a neighbor at both sides. Therefore, for a population of individuals on a ring or circle, the coalescent probabilities in a single generation follow a uniform distribution:

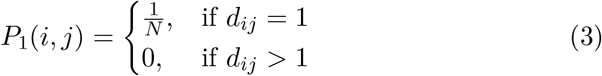

where *d_ij_* is the linear distance between *a_i_, a_j_*. On a bounded line, the probability of adjacent pairs coalescing in the previous generation is *P_1_*(*i, j*) = 1/(*N* - 1).

One of the consequences of structure in this one-dimensional model is a spatial separation of lineages in the genealogy, such that a haplotype *a_i_* with an ancestor *a_i_*(*t*) at position *i* – 1 in generation *t* is prevented from simultaneously having an ancestor at position *i* + 1 at generation *t* + 1. In forward time, this separation of lineages occurs because an individual at position *i* + 1 that was replaced by the offspring of its right neighbor cannot simultaneously be the grandchild of its left neighbor, because an individual at location *i* + 1 cannot be the parent of one at *i* – 1, or the reverse. This constraint is shown schematically in Figure 1. As is the case for the coalescent in an island model, we predict that the consequence of lineage separation will be a tendency for individuals sharing a more recent common ancestry to cluster together spatially, and to increase the expected coalescent time (branch lengths) for a pair of individuals that are selected at random with respect to their location.

**Figure 1.**
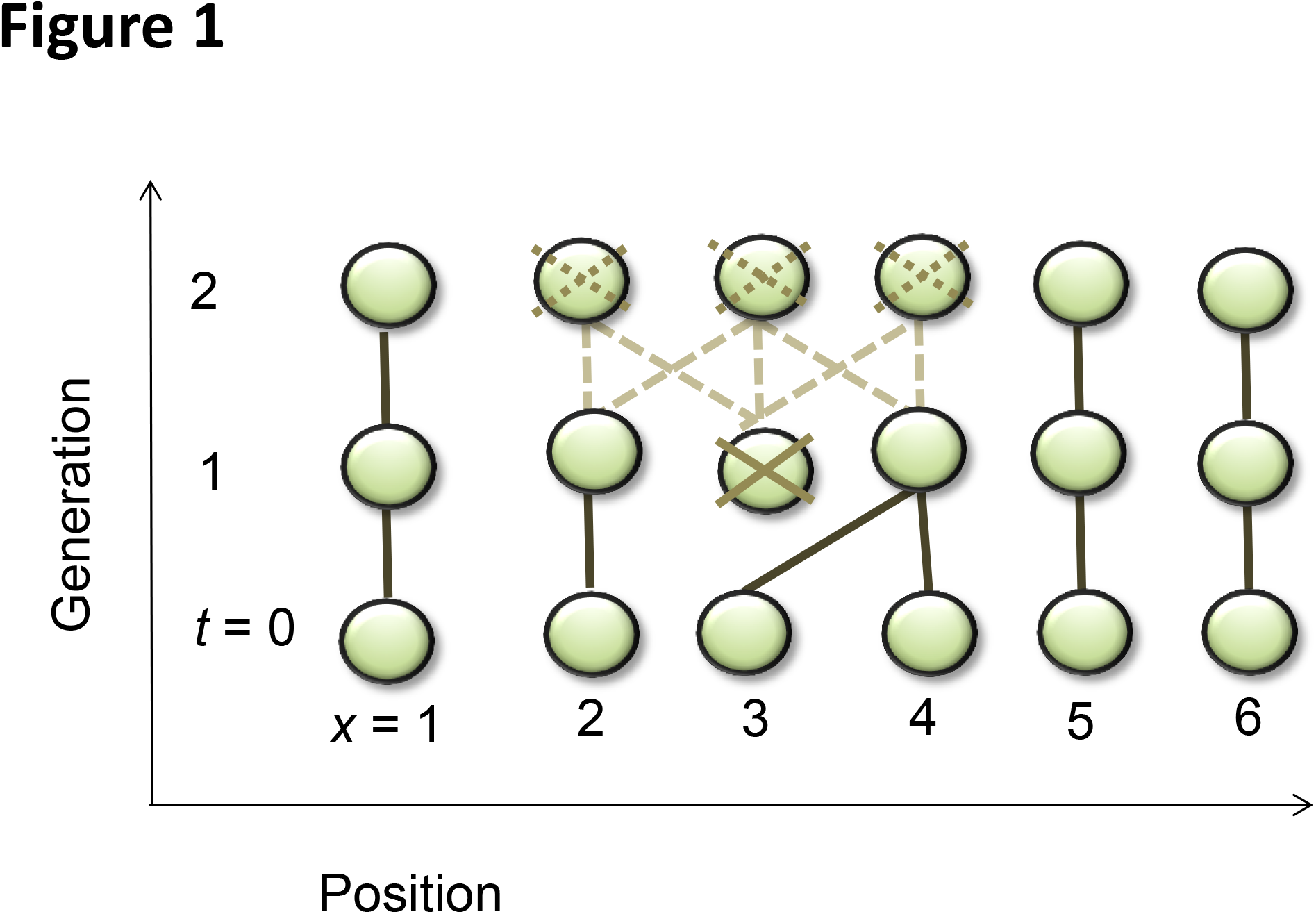
The death of the individual (haplotype) at position 2 at in generation *t* = 1 leads to a separation between the lineages of *a_2_* and *a_4_*. Specifically, none of the possible deaths and replacement events among individuals at positions 2,3,4 between generations *t* = 1 and *t* = 2 (shown as dashed lines) corresponds to a co-ancestry between *a_2_* and *a_4_* at *t =* 2.

It also follows that the probability of either *a_i_* being a descendant of an individual at position *j* or vice-versa is *P*_1_(*a_i_ → j*) = *P*_1_ (*a_j_ → i*) = 1/2*N* (by symmetry, half of Eqn. 3), where *a_i_ → j* denotes *a_i_* having an ancestor in position *j*. Using these coalescent probabilities, we can define a matrix of transition probabilities **M**; the matrix elements *m_ij_* are the probabilities that each haplotype *i* is descended from an “ancestor” (including the possibility of persistence) in position *j* in the previous generation. The off-diagonal coefficients are then the probabilities of individuals *i* and *j* coalescing in the previous generation. The diagonal coefficients are the probabilities of an individual persisting without death or replacement in previous generations at the same spatial position, i.e. *a_i_ → i*.

For a population on a circle, where there are no boundary asymmetries, the probability that haplotype *j* is descended from *i* in the previous generation is *m_ij_*, represented by symmetric transition matrix:

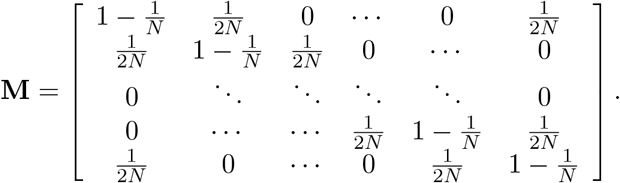

In a ring-shaped population structure, the ordering of points (i.e. which is labeled 1,2 etc) is arbitrary. In contrast, a population on a bounded line introduces the boundaries with unidirectional coalescent paths at the endpoints, the transition matrix **M** is asymmetric,

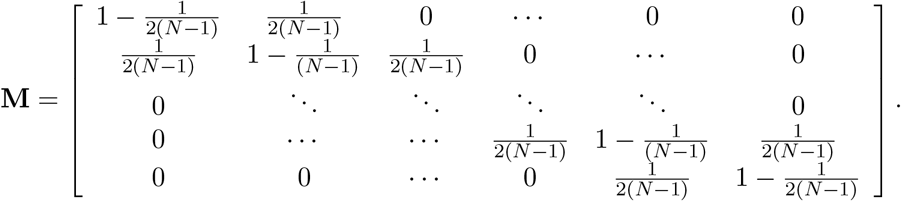

The structure of these transition matrices is similar to that of stepping stone island models of migration (e.g. Maruyama (1970), Nagylaki (1974), Slatkin (1991, 1993)), for the special case of deme size *n* = 1 and with the per-generation probabilities of migration replaced by coalescent probabilities. Because M provides a dynamically sufficient description of the coalescent processes in circular and linear populations, the expected time to a coalescent event for any pair of *i, j* can be computed from these matrices as a first passage time between the two states (i.e. for both *i* to *j* paths and vice-versa).

To define the distribution of coalescent probabilities in terms of a Markov chain, let *X_t_,Y_t_* be the positions of the lineages of individuals *a_i_*(*t*), *a_j_*(*t*). The probability of a coalescent event at time *t* is:

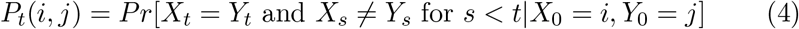

This is equivalent to defining a single Markov chain on bivariate *Z_t_* = (*X_t_,Y_t_*). The transition probabilities for this bivariate Markov chain are defined by the products of transition matrices on the individual lineages

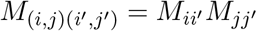

for any *i ≠ i’, j ≠ j’*. Defining a bivariate diagonal *D* in the space of *Z, D* = (*k, k*), *k* ∈ (1…*n*), being in the set of *D* corresponds to the lineages of *a_i_, a_j_* coalescing at some *k*. The probability of coalescing at time *t* is then

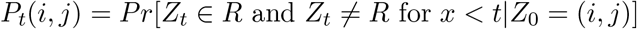

These transition probabilities can be computed recursively using the following equation:

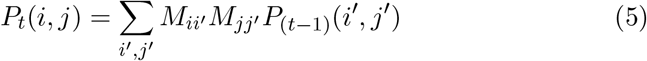

for *i ≠ j* and *t* > 0, with initial conditions *P*_0_(*i, j*) = 0 for *i ≠ j*, 1 for *i = j*.

We will apply this recursion to compute coalescent probabilities and coalescent times for the one dimensional spatial model by leveraging the symmetries of the transition matrices, specifically the equivalence relation among pairs characterized by distance *d_ij_* between *a_i_*(0),*a_j_*(0) on a bounded line or on a circle. Using the spatial distance 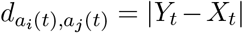 will allow us to reduce the general bivariate Markov chain described above to a simpler univariate model.

In a single generation, the transition among haplotype pairs *a_i_*(0), *a_j_*(0) → *a_i_*(1), *a_j_*(1) maps to a transition in their spatial distance 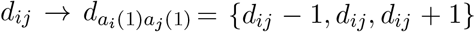. These respective distances correspond to the following outcomes: in the first instance, the previous generation’s ancestors *a_i_*(1), *a_j_*(1) of *a_i_*(0), *a_j_*(0) are located one unit closer to one another than their initial distance *d_ij_*. The second case occurs in the absence of a coalescent event involving *a_i_* or *a_j_*, while the third case, an increase in distance, occurs when either *a_i_* or *a_j_* coalesce with neighbors in a direction opposite to one another.

For two haplotypes located on a circle that are not at maximum distance *d_max_* = ⌊*N*/2⌋, the transition probabilities *∏* for the ancestral distances 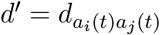 in a single generation are

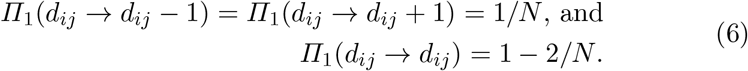

When the haplotypes are located internally on a bounded line, the transition probabilities are the same as above apart from having *N* – 1 in place of *N*.

Asymmetries in these transition probabilities occur when one or more of the haplotypes in a pair is an endpoint on a bounded line, or if the distances between haplotypes on a circle is the maximum value *d_max_* = ⌊*N*/2⌋. For the endpoints of a bounded line, coalescent events can only occur in a single direction, while for two points *i, j* at *d_ij_ = d_max_* on a circle, any coalescent events involving either *i* or *j* will either bring *a_i_* and *a_j_* closer together (if *N* is even) or maintain the same distance (if *N* is odd), while coalescent events in relation to other haplotype pairss can be in either a *d_ij_* + 1 or *d_ij_* – 1 direction. These alternative scenarios are illustrated in Figure 2, and the transition probabilities corresponding to different points on the circle and on the line are summarized in Tables 1–2. The asymmetries caused by these edge and midpoint cases are negligible for large population sizes because of the much larger number of transition probabilities that are identical, as are the differences due to the 1/(*N* – 1) vs. 1/*N* transition probabilities for circular vs. bounded linear spatial structures (Eqn. 3).

**Figure 2.**
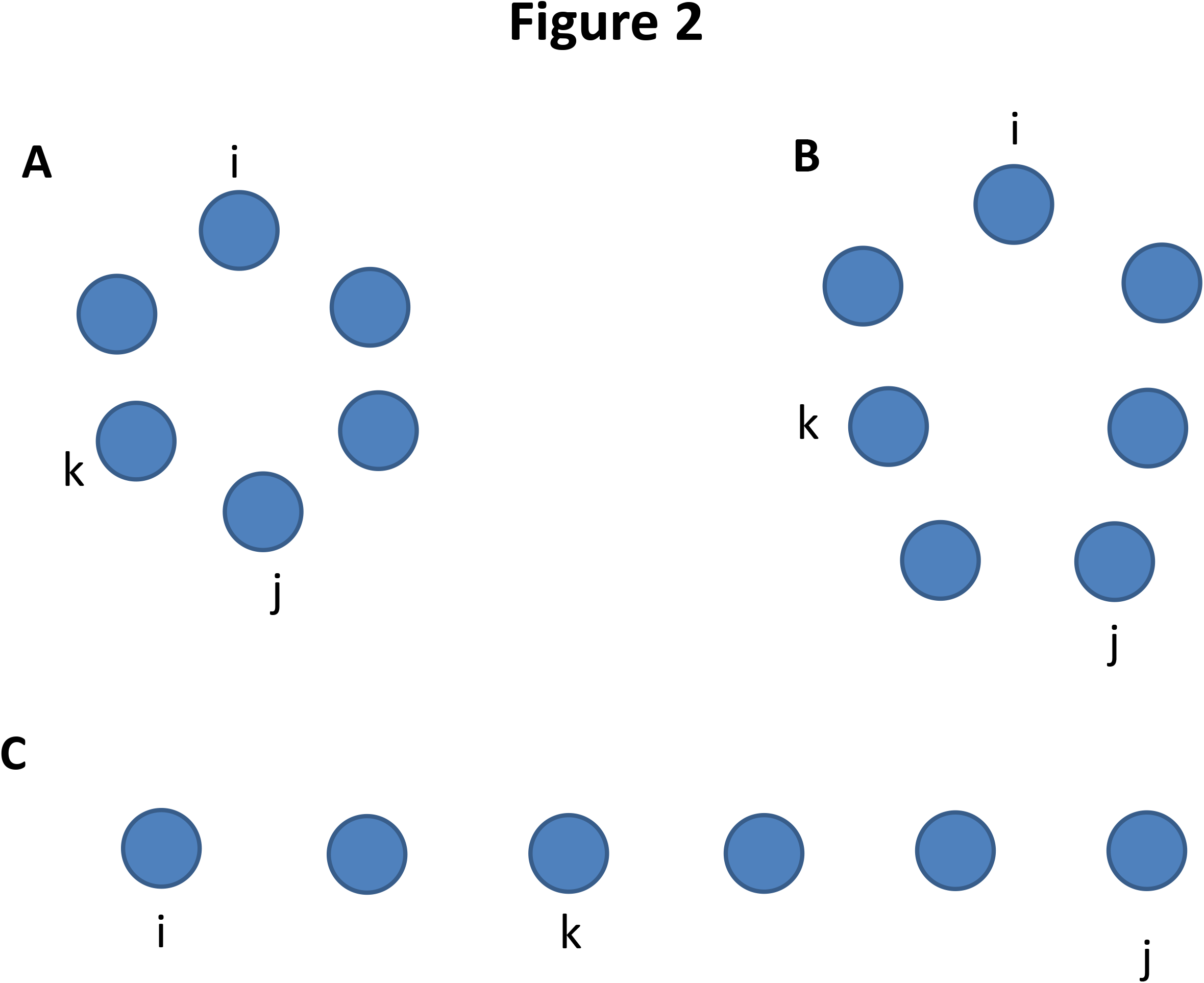
Scenarios of one-dimensional spatial structure considered in this paper, and the possible positions/distances that lead to asymmetries and boundary effects in transition probabilities. A) a circular population with an even *N* = 6. Individuals *i, j* are at *d_max_* = 3, so that any coalescent event of *i* or *j* with a neighbor *k* reduces the ancestral distance. The same is not true in B. with an odd (*N* =) number of entries, where a coalescent of *i* towards *k* reduces the distance while *j* towards *k* decreases it. Figure 2C shows a bounded linear population structure, where endpoints *i, j* lead to asymmetry due to boundary conditions, both with respect to mutual distance transition probabilities and with respect to internal point *k*.

**Table 1.**

| **Transition** | **Interior** | **$N$ even, $d = N/2$** | **$N$ odd, $d = (N - 1)/2$** |
| --- | --- | --- | --- |
| $d \rightarrow d - 1$ | $\frac{1}{N}$ | $\frac{2}{N}$ | $\frac{1}{N}$ |
| $d \rightarrow d + 1$ | $\frac{1}{N}$ | 0 | 0 |
| $d \rightarrow d$ | $1 - \frac{2}{N}$ | $1 - \frac{2}{N}$ | $1 - \frac{1}{N}$ |

In a population of *N* haplotypes on a bounded line, the transition probabilities of the pairwise distances are given in Table 2,

**Table 2.**
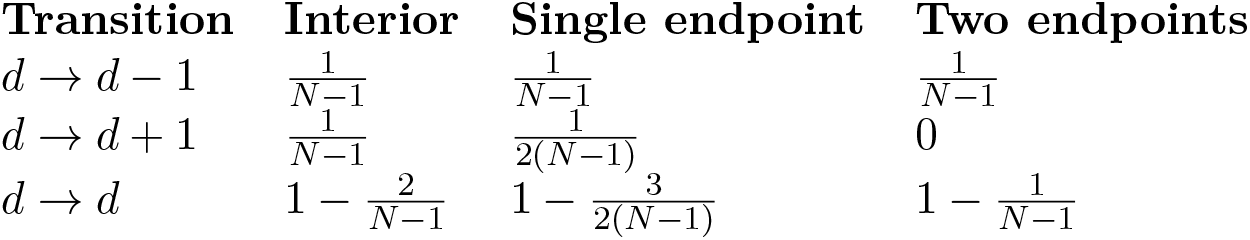

To compute the coalescent probabilities of a randomly selected haplotype pair, we would like to specify the transition probabilities conditional on distance (with *d_ij_* = 1 as a base case) and the distribution of pairwise distances over all possible pairs. There are (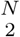) such pairs in a population of size *N*. In a circular population, there are *N* pairs for every possible distance *d_ij_* = 1, 2…⌊*N*/2⌋, so that the probability of a randomly selected pair being at distance *d* on the circle is

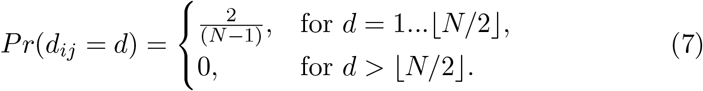

Among pairs of haplotypes on a bounded line, there are *N — d* possible pairings at distance *d* due to the absence of “wraparound” pairings, so that the distribution of pairwise distances is

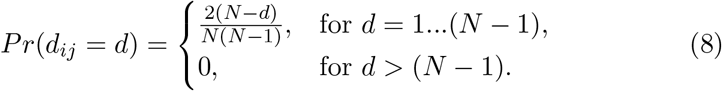

Using the coalescent probabilities in a single generation (Eqn. 3), the transition probabilities on distances (Eqn. 6, Tables 1–2), and the distribution of pairwise distances in Eqns. 7–8, we construct a system of equations for the probabilities of a random pair at positions *i, j* coalescing in generation *t* generation

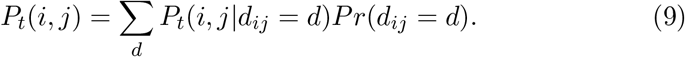

The transition probabilities *P_t_*(*i, j*|*d*) can be computed recursively as a sum over all possible ancestral distances *d’ = d — 1, d, d* + 1 multiplied by the base case transition probabilities for a single generation:

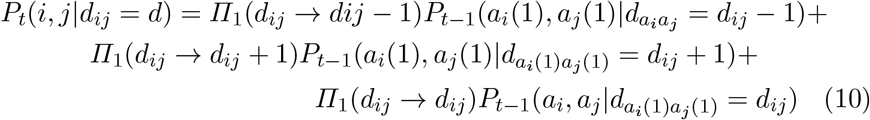

where *∏_t_(d → d’)* is the probability that haplotypes *a_i_, a_j_* at distance *d = d_ij_* have ancestors *a_i_*(*t*), *a_j_*(*t*) at distance 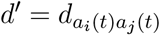, and where by definition *d* = 0 corresponds to a coalescent event. This recursive relationship is a special case of Eqn. 5, a consequence of the equivalence among all pairs *i, j* with a spatial distance *d_ij_* = |*X_t_* – *Y_t_*|.

In order to develop an intuition for the effect of spatial separation of lineages in this one-dimensional model on the Moran coalescent, we compare coalescent probabilities for *t* = 1, 2 generations on the circle to the corresponding values in the unstructured Moran model. In the absence of spatial structure, the coalescent probability of the pair *a_i_, aĳ* in a single generation in the Moran model is the probability of picking that particular pair from all possible pairs

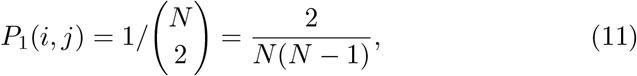

while the coalescent probability in *t* generations is

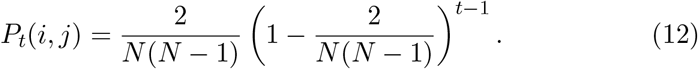

In contrast, for a population on a circle, the coalescent probability in a single generation is

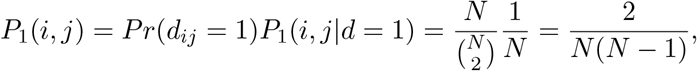

which is identical to the single generation coalescent probability in a Moran model without spatial structure. If the population is linear rather than circular, there is a slight difference due to edge effects that introduces a factor of *N*/(*N* – 1), which is effectively ≈ 1 when *N* is large.

The effect of population structure becomes apparent for *t* ≥ 2 generations, i.e.

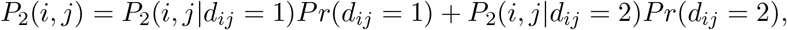

which follows because *P_2_*(*i, j*|*d_ij_ > 2*) = 0. From Eqn. 7, the probability of selecting a pair at *d* = 2 for a circular population is

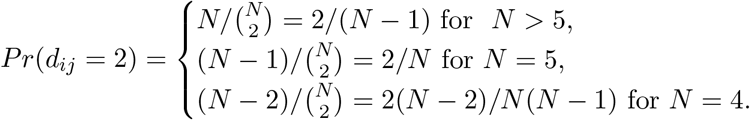

while in a bounded linear population with *N* > 3 there are *N* – 2 possible *d* = 2 neighbors, so that 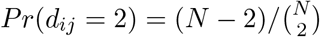.

To expand the recursion in Eqns. 9–10 for *t* = 2 generations, we can equate the probability that *a_i_, a_j_* at distance *d_ij_* = 1 coalesce in 2 generations to the product of probabilities that they do not coalesce with one another in *t* = 1 followed by the probability of coalescing at *t* = 2. Without loss of generality, assume that *i* is located to the left of *j*. Because of the directional separation of lineages shown in Figure 1 in order for *i, j* to coalesce at *t* = 2, we exclude coalescent events where *a_i_*(1) is to the left (or counterclockwise) of *a_i_* (i.e. an individual at position *i* replaced by a descendant of its left neighbor), as well as cases where *a_j_*(1) is to the right of *a_j_*, because the lineages of *a_i_* and *a_j_* are spatially incapable of coalescing at *t* = 2 under these scenarios. In contrast, if a_i_(1) is to the right of *a_i_* or *a_j_*(1) is to the left of *a_j_*, the lineages are not spatially separated and can coalesce in two generations. In forward time, this corresponds to a “grandchild” of a haplotype at position *i* replacing another at *j* or vice-versa.

The probability of neighbors coalescing in exactly 2 generations is then:

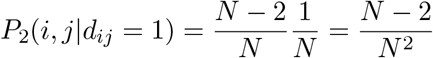

The factor *N*–2 in the numerator of the first term of the sum is the combined probability that neither *i* nor *j* coalesce with one another at *t* = 1, nor coalesce with their left or right neighbors through replacement, whereby the latter event “prevents” coalescing at *t* = 2.

If *d_ij_* = 2, the only means by which a coalescent event can occur in two generations is if either *i* or *j* coalesce with *k* located between *i* and *j* at *t* = 1, such that *k* is the ancestor of either *i* or *j*, followed by a coalescent event between *k* and the remaining haplotype at *t* = 2, i.e.

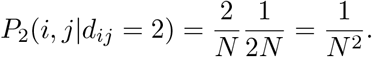

The exception to this transition probability is for the special case of *N* = 4, where there are two possible paths for *d_ij_* = 2 pairs (because if *d_ij_* = 2, both the left and right neighbors are mutually *d* = 1 with respect to both *i* and *j*), so that 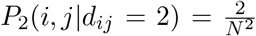. Excluding these special cases, following Eqn. 13, the probability of coalescing in two generations when *N* > 5 is

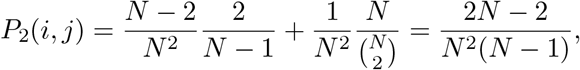

while for the exceptional cases in which the population size *N* < 6 the coalescent probabilities are:

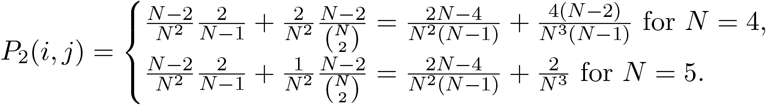

These probabilities are smaller than the probability of coalescing in two generations in the unstructured Moran model, leading to longer terminal branch lengths (expected coalescent times) in the genealogies. Computing from the weighted sum over coalescent probabilities over all possible coalescent times, the expected coalescent time *t* for a sample pair is:

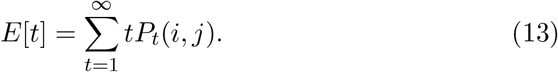

Similarly, the variance and all higher moments *t^k^* of the distribution of coalescent times are given by:

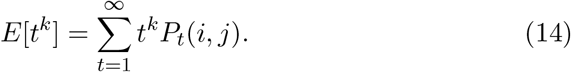

For comparison, the expected coalescent time *t_m_* in the Moran model is

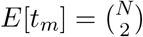

and the variance in coalescent time is

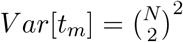

In the following section, we use Eqs. 9-10 to calculate the expected coalescent times on a circular population structure for several population sizes. These numerical estimates of coalescent probabilities and coalescent times are compared to those observed in individual-based simulations and to estimated coalescent times and prbabilities in the unstructured Moran model.

## 3 Numerical and Simulation Results

Our simulations of the Moran coalescent in one spatial dimension will focus on circular population structure, because the model’s symmetry makes comparison to numerical predictions more straightforward, and because populations with bounded linear structure will show qualitatively similar spatial effects except when population sizes are very small. We also only consider even population sizes without loss of generality, as the contribution of the difference in transition probabilities on circle midpoints (Table 1) to the overall distribution of pairwise distances will be trivial unless *N* is extremely small.

For the set of population sizes *N* = {10, 20, 50, and 100}, we simulated Moran genetic drift with one dimensional spatial structure by selecting a haplotype at a random position to die in every generation. An individual at this random position *i* is replaced with equal probability by the progeny of either the haplotype at position *i* + 1 or at *i* – 1. In cases where *i* is located at an endpoint, the left or right neighbor is replaced by those at positions 1 or *N* to be consistent with probabilities in an unbounded (circular) population. Ancestor-descendant relations from the simulated genealogies were stored in a tree data structure to facilitate tracing back the genealogy of the population. The coalescent probabilities for each time unit were estimated over 1000 (*N* = 10, 20) and 100 (*N* = 50, 100) replicates, respectively by counting over coalescent events at each *t* over all replicates. Coalescent times were computed over all pairs across simulations using traceback to the most recent common ancestor. The distributions of coalescent times and probabilities in the simulations are compared to those obtained numerically from the recursions in Eqn. 10. All code was implemented using Python 2.7.3 and is available from the corresponding author upon request.

In the absence of population structure, the pairwise coalescent probabilities follow a geometric distribution (Eqn. 12), so that the expected coalescent time for a pair of haplotypes in a population of *N* is *N*(*N*–1)/2 ~ *N*^2^/2.

It has also been shown (Moran, 1958, Moran and Watterson 1959) that the expected coalescent time for all *N* haplotypes in an unstructured Moran model is ~ *N*^2^ (Eqns. 13–14). Consequently, all of our simulations were run for ~ *N*^2^ generations or more when computationally feasible, to ensure the coalescence of all haplotypes in the population. When computationally feasible (i.e. for smaller population sizes) these extended simulations were run *t* ≫ *N*^2^ because the structured models has greater expected coalescent times among haplotype pairs than in the standard Moran model.

Figures 3A-C compare coalescent probabilities in the structured model (in both simulations and the recursion) to the unstructured Moran coalescent, while the boxplots in Figures 4a-d compare the distributions of coalescent times computed from the recursions for different population sizes. As predicted analytically, the coalescent probabilities in the structured model are smaller than in the unstructured model for time intervals *t* ≪ *N*^2^ (except *t* = 1 where they are equal), as a consequence of the spatial separation of lineages. As *t* approaches *N*^2^, the values of *p_t_* in the structured and unstructured models eventually cross so that the coalescent probabilities are actually greater in the structured model for sufficiently large *t* than those in the unstructured Moran model. This is because most of the coalescent events occur earlier in the unstructured model, while due to lineage separation, some haplotypes remain uncoalesced at high *t* in the spatially structrued population. This can be seen most clearly from the intersection of estimated *P_t_* in the structured and unstructured models in Figures 3A-B (the same would be the case in Figs. 3C-D if much longer time intervals were represented for these larger population sizes).

**Figure 3.**
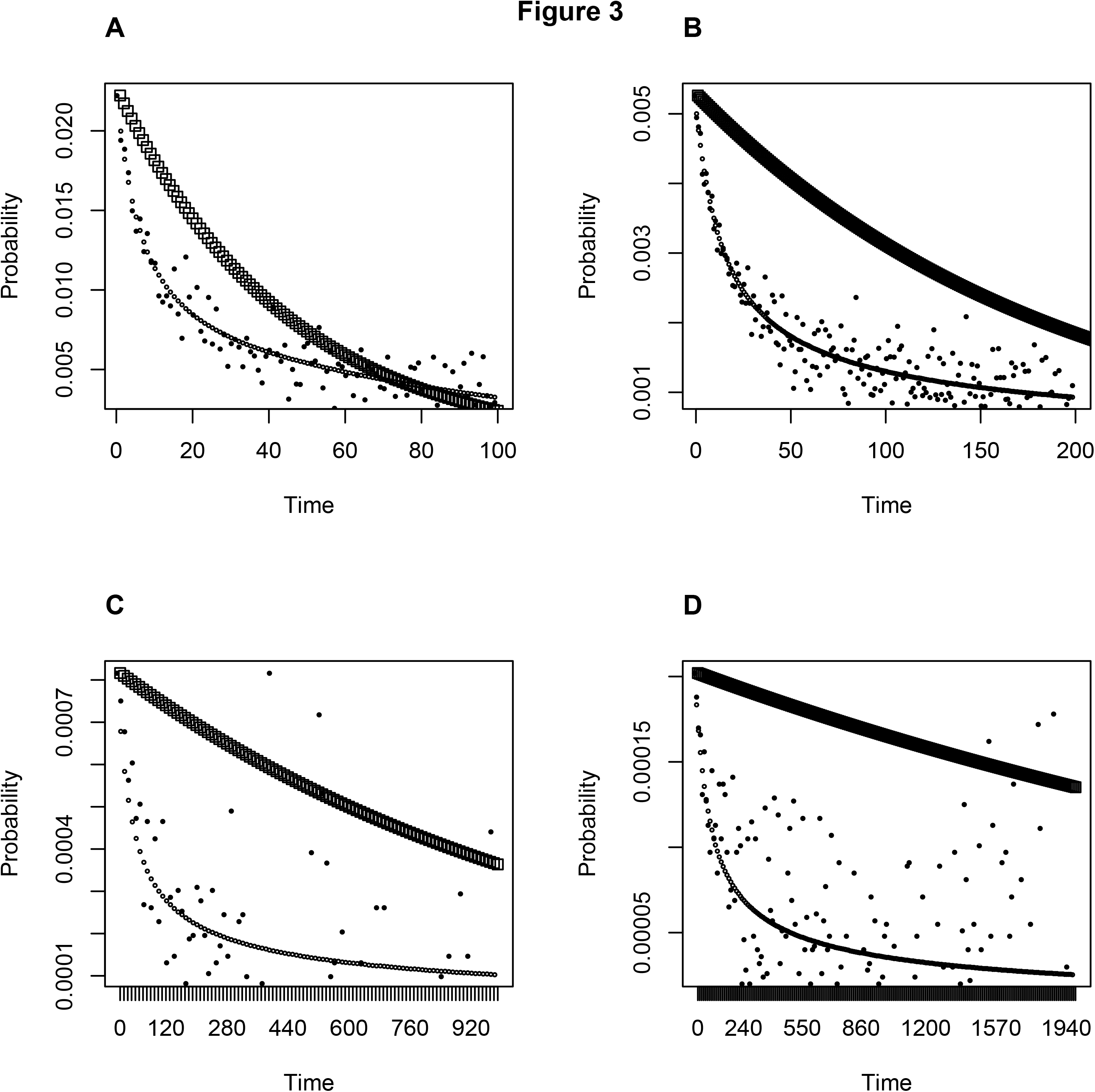
Numerical estimates of coalescent probabilities as a function of time for the circular structured model (open circles) and unstructured Moran coalescent (open squares) plotted against coalescent properties estimated from individual-based simulations of the Moran process in a circular structured population (solid circles). A) population size *N* = 10, B) *N* = 20, C) *N* = 50, and C) *N* = 100. For Figures 3C and 3D, time points were sampled in 10 and 20 unit intervals, respectively, in order to provide a legible density of data points comparable to those in Figures 3A-B. Note the close fit between simulated and numerical values for higher probabilities associated with small *t* and lower *N*. The intersection of coalescent probabilities in the structured vs. unstructured models can be seen in Figures 3a-b, the trajectories cross eventually for larger population sizes as well for *t* ≫ *N*^2^, i.e. longer time intervals than those sampled in the simulations.

**Figure 4.**
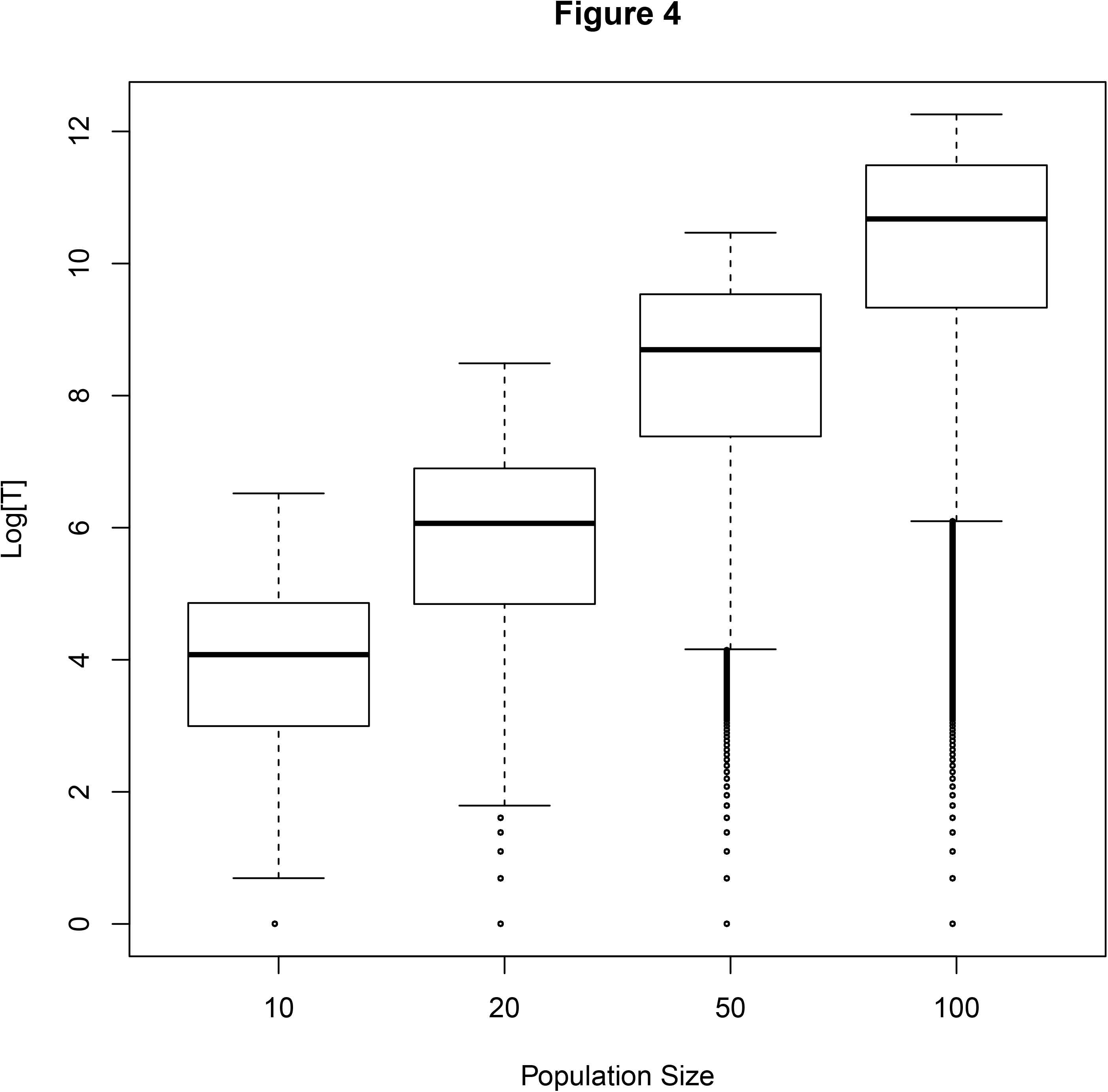
Boxplots showing the distribution of coalescent times (on a logarithmic scale) among pairs for populations of sizes A. *N* = 100, B. *N* = 20, C. *N* = 50 and D. *N =* 100 computed numerically and pooled over 1000 (A-B) and 100 (C-D) replicates for consistency with the simulations. The mean values are the *E[T*] values in Table 3, while the spread of the quartiles is consistent with the magnitude of the standard deviations from Table 3, i.e. *Var[T] ~ E[T*]^2^.

The eventual coalescence of most lineages also explains the good fit of *P_t_* estimated from simulations to the numerical estimates when *t* is small and when coalescent probabilities are relatively high. For small *t*, most deaths and replacements correspond to the most recent coalescent event for some pair of haplotypes, so that in any sample path the number of coalescent events matches the expected number at *t*. As *t* increases and more lineages coalesce, many death and replacement events involve pairs of lineages (defined at *t* = 0) that have already coalesced, while others still correspond to most recent common ancestors. As a result, there is a large variance in whether a coalescent event occurs for a particular sample path *t*. This variance scales with the fraction of lineages that has coalesced, which can be orders of magnitude larger than *P_t_* itself. Specifically, the fraction of pairs that have coalesced at time *T* is 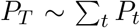 so that the Bernoulli variance in whether a birth/death event corresponds to a most recent common ancestry scales as ~ *P*_T_(1 — *P_T_*)/*R* for *R* replicate sample paths. Consequently, in any sample path there will always be either more or fewer coalescent events at any large *t* than the predicted value. This is particularly pronounced in Figures 3C-D for *N* = 50, 100, because the coalescent probabilities for larger *t* are of the order ~ 10^-^5 – 10^-^4, and are estimated over only *R* = 100 replicates. This accounts for the large scatter of estimated coalescent probabilities in Figures 3C and 3D when *t* is large.

The observed differences between coalescent probabilities for a fixed *t* between the structured and unstructured models may seem trivial, but they reflect qualitatively significant differences in the shape of respective genealogies. Namely, the deeper internal branch lengths in the structured model are associated with much longer coalescent times, particularly for populations that are large enough for multiple subsets of haplotypes to achieve spatial separation over many generations. This is a consequence of the fact that two haplotypes (and their ancestral lineages) at a distance *d_ij_* have to wait a minimum number of generations equal to their distance of separation before a coalescent event can occur (and for large d, the typical coalescent time will be much more than *d* generations). The expected coalescent times and their variances in the structured Moran model are shown for *N* = 10, 20, 50, 100 in Table 3 and compared to those of the unstructured model. Even for modest population sizes of *N* = 50,100 the expected coalescent times are an order of magnitude greater with spatial structure (*E*[*T_s_*], and its variance *Var*[*T_s_*]) than in its absence (*E*[*T_m_*]).

**Table 3.**

| $N$ | $E[T_s]$ | $Var[T_s]$ | $E[T_m]$ | $Var[T_m]$ |
| --- | --- | --- | --- | --- |
| $N = 10$ | 91.67 | 1022.22 | 45 | 2025 |
| $N = 20$ | 700 | 632100 | 190 | 36100 |
| $N = 50$ | 10625 | $1.228 \times 10^8$ | 1225 | $1.501 \times 10^6$ |
| $N = 100$ | 84120 | $9.707 \times 10^9$ | 4950 | $2.450 \times 10^7$ |

The distributions of coalescent times are shown as boxplots in Figures 4A-D for numerical estimates calculated from the recursions. Both the mean values (sample estimates of *E*[*T*]) and the magnitudes of the outer quartiles are consistent with the first and second moments computed numerically from the recursions in Eqs. 9-10.

## 4 Discussion and Future Directions

The strict spatial constraint of only allowing neighbors to coalesce imposes changes to the distribution of coalescent probabilities and the shape of the coalescent genealogy. These changes have the potential to distort our estimates of population parameters and our inferences of evolutionary and demographic processes. As was noted in the introduction, the pairwise genetic distance 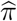 is the Tajima (1989) estimator for the population mutation rate *θ* = 4*Nµ*. Because 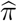 scales linearly with the mean coalescent time among haplotypes in a population (Eqn. 1), the inflation of *E[T]* by orders of magnitude due to the spatial constraint will similarly inflate our estimates of *θ*. The practical implications of this are that population mutation rate *θ* and effective size *N_e_* (when mutation rates *µ* are known approximately) will tend to be substantially over-estimated for random samples of haplotype from spatially structured populations.

As a consequence, the increase in estimates of *θ* due to longer coalescent times (Table 2) can result in false inferences of natural selection or of recent changes in population size. The prevalence of longer internal branch lengths among spatially distant haplotypes correspond to patterns mimicking the distribution of branch lengths observed under diversifying selection, as well as the shape of coalescent trees often seen following recent population bottlenecks where only the most common variant alleles (corresponding to the deepest internal branches) persist. Conversely, because this constrained linear structure inflates 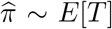, it can also obscure recent signatures of purifying selections or population expansions, because the increase in genetic distances due to structure act as a counterweight to the reduction in 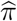 due to purifying selection.

The implications of such results are especially pertinent to population genetic studies of organisms with limited motility and dispersal. These include vegetatively reproducing plants or those with non-dispersing seeds, sessile marine invertebrates with a non-planktonic egg and larval state, and particularly colonies of microbes and other single-celled organisms. In cancerous tumors, cells belonging to related subclonal lineages tend to occur in close spatial proximity. The direct applicability of a one-dimensional model of spatial structure may be somewhat limited, since relatively few organisms are spatially constrained onto a one-dimensional string. At a microbial scale, there are linear or near-linear population structures (e.g. many filamentous cyanobacteria and Streptococcal bacteria, and the arrangement of epithelial cells in relation to stem cells in colorectal crypts is linear as well), and at a more macroscopic scale, low-dispersal species that are tightly restricted to ridgelines or riparian and tidal zone environments might adhere somewhat to this linearly constrained model. Moreover, we expect that some qualitatively similar effects will be found in spatial models extending into two or three spatial dimensions. A two-dimensional lattice may be a good approximation to the coalescent in microbial biofilms, while three dimensional lattices would provide a closer model for the population structure of spherical colonial algae (e.g. *Volvox*), Staphylococcal clusters, and among subclonal cell lines in cancerous tumors.

Based on the qualitative interpretation of our findings from a spatially constrained linear coalescent, it would follow that analyses of genetic variation that compare single nucleotide variant or haplotype frequency distributions in tumors to standard neutral models could lead to false inferences of selection. Specifically, when all samples come from a small sector, genetic variation and *θ* will be underestimated while samples from distant sectors will lead to overestimates (e.g. Shpak and Lu 2016). Researchers such as Ling et al (2015) have dealt with this issue by sampling multiple sectors from different regions of large tumors, while Waclaw et al (2015) explicitly simulated mutation and drift in a three dimensional spatial model. In order to develop an analytical framework for neutral evolution in tumors as a proper null model for detecting selection, distributions of coalescent times for three dimensional spatial models may be required. Ideally, coalescent times and probabilities could be computed using the Euclidean distance between haplotypes as a dynamically sufficient characterization (as for this model), and integrated over all possible distances to obtain the coalescent probabilities.

The principal difference between the linear model and higher-dimension models is the greater number of degrees of freedom for each coalescent event in multiple dimensions. Specifically, the number of possible *d* = 1…*k* neighbors and the number of possible coalescent paths between any two haplotypes in more than a single dimension will depend upon how closely packed individuals are and on the specific geometry of the packing. Presumably, the larger number of neighbors for each individual creates more mixing of the population in comparison to the linear model and consequently less prounounced increases to the expected coalescent times, although this remains to be demonstrated in the general case.

Generalizing the recursions presented here to such cases represents an important research question because of the broad applicability of the spatially structured coalescent to many natural and model populations (see for example Kelleher et al 2014 for simulations of coalescent processes on multidimensional grids). The approach taken in this paper has the advantage of parameterizing coalescent probabilities solely as functions of spatial distance rather than in terms of transition probabilities among specific pairs, i.e. treating all pairs with the same distance as equivalence classes, this is an approach with the potential be generalized to higher-dimensional models, thereby reducing their intrinsic computational complexity and the number of state variables.

## 5 Acknowledgements

We thank Jon Wilkins, Mark Kirkpatrick, Peter Mueller, and Gordon Zitkovic for helpful discussion. MS and JL were supported by funding from the St. David’s Foundation Impact Fund in 2016. JPT was supported by the Notsew Orm Sands Foundation.

